# Noise-Aware Constitutive Compatibility Atlas for Hyperelastic Material Characterization

**DOI:** 10.64898/2026.10.12.758616

**Authors:** Amirreza Asadi, Kaveh Laksari

## Abstract

Hyperelastic constitutive characterization is usually posed as selecting the best-fitting model, although several families can reproduce measured loading paths while predicting different responses elsewhere. We developed a noise-aware constitutive-response atlas to determine which families are compatible with an observed response over a prescribed deformation range and within a stated error tolerance. Each family forms a continuous response manifold in a common bounded deformation space, with exact nesting preserved. Standard uniaxial, biaxial, and shear tests are restricted projections of the complete response space, allowing the atlas to identify deformation states that resolve ambiguities left by conventional calibration. The output can contain several families, one family, or none, without forcing a single winner. We constructed the atlas for seven incompressible isotropic hyperelastic families using literature-based parameter ranges. Numerical certification and three controlled tests examined generating-family recovery, separation of responses distinct at the prescribed tolerance, and rejection of a withheld constitutive law. External validation used Treloar rubber and human-brain datasets. For Treloar rubber, uniaxial calibration retained the Mooney-Rivlin and one-term Ogden families; an atlas-selected equibiaxial measurement favored Mooney-Rivlin. For cortex, compression retained the one-term Ogden and second-order Yeoh families; withheld tension and shear favored one-term Ogden. The atlas thus preserves genuine ambiguity, exposes an inadequate candidate library, and identifies when another measurement is needed.

## 1. Introduction

Many hyperelastic constitutive families are used to represent soft biological tissues as well as many rubber-like materials. Several constitutive families can produce identical measured loading paths but diverge elsewhere. For this reason, characterization has moved beyond least-squares error toward likelihood-based comparison, Bayesian calibration and evidence, and mechanics-informed classification [1–5]. These approaches quantify parameter uncertainty and compare fitted families, but their conclusions still depend on which families are considered and which deformations are observed. That choice can materially affect subsequent stress, strain, and biomechanical predictions [6–9].

A low fitting error does not imply unique constitutive identification. Distinct families may be indistinguishable along restricted experimental paths [3,10], and Bayesian studies likewise show that fit quality alone cannot establish adequacy [4]. Some ambiguity is structural rather than numerical: lower-order constitutive laws are exact subsets of richer families. A sound characterization should retain every family supported by the data and should also allow the possibility that none of the configured families is adequate.

Material Fingerprinting takes a related approach: measured responses are compared with precomputed fingerprints, avoiding repeated continuous optimization [11,12]. Adaptive extensions build increasingly expressive combinations of admissible hyperelastic features [13]. From a geometric viewpoint, the responses generated by one family form a lower-dimensional manifold in observation space [14,15]. We use this view to ask which family manifolds intersect a prescribed tolerance neighborhood around the observation. The aim is not to return only the nearest fingerprint or to force one family to win.

Material Fingerprinting and the present atlas share a practical motivation, i.e., replacing costly repeated optimizations with a precomputed lookup library, but they tackle different questions. While Material Fingerprinting reports the single admissible instance and material family whose precomputed response lies closest to a measured observation [11–13], the compatibility atlas indicates whether a family’s response manifold intersects a predefined tolerance neighborhood around the observed values, which could contain multiple families. This distinction has three consequences. First, when several manifolds pass through the same tolerance neighborhood, which as we will see occurs for a substantial fraction of the constructed atlas, compatibility reports the full retained set rather than an arbitrary nearest choice, which could be chosen due to numerical considerations rather than physical attributes. Second, exact nesting between families is enforced through hierarchy closure, so a compatibility label reflects genuine mathematical containment rather than numerical proximity. Third, when no family in the library falls within the required tolerance, the atlas returns an empty set rather than the least-bad match; nearest-fingerprint frameworks have no analogous mechanism for signaling that the candidate library itself is inadequate.

Set-membership identification and bounded-error validation treat consistency as a feasibility question, not simply as a ranking problem [16,17]. We apply the same principle to competing nonlinear hyperelastic families represented in a common deformation-response space. Uniaxial (UT), equibiaxial (BT), and simple shear (SH) are treated as restricted probe paths through the admissible multiaxial deformation space. Instances from different families can agree along those paths and separate at other admissible states [8,9,18,19].

We therefore formulate hyperelastic characterization as a family-level compatibility problem. The atlas combines continuous response manifolds, a declared discrepancy tolerance, and exact hierarchy closure. We construct and numerically certify it for seven incompressible isotropic families, examine three controlled cases, and test it prospectively with withheld loading modes from Treloar rubber and human brain tissue. The reported outcome is the adequate family set, which may contain several families, one family, or none (Figure 1).

**Figure 1.**
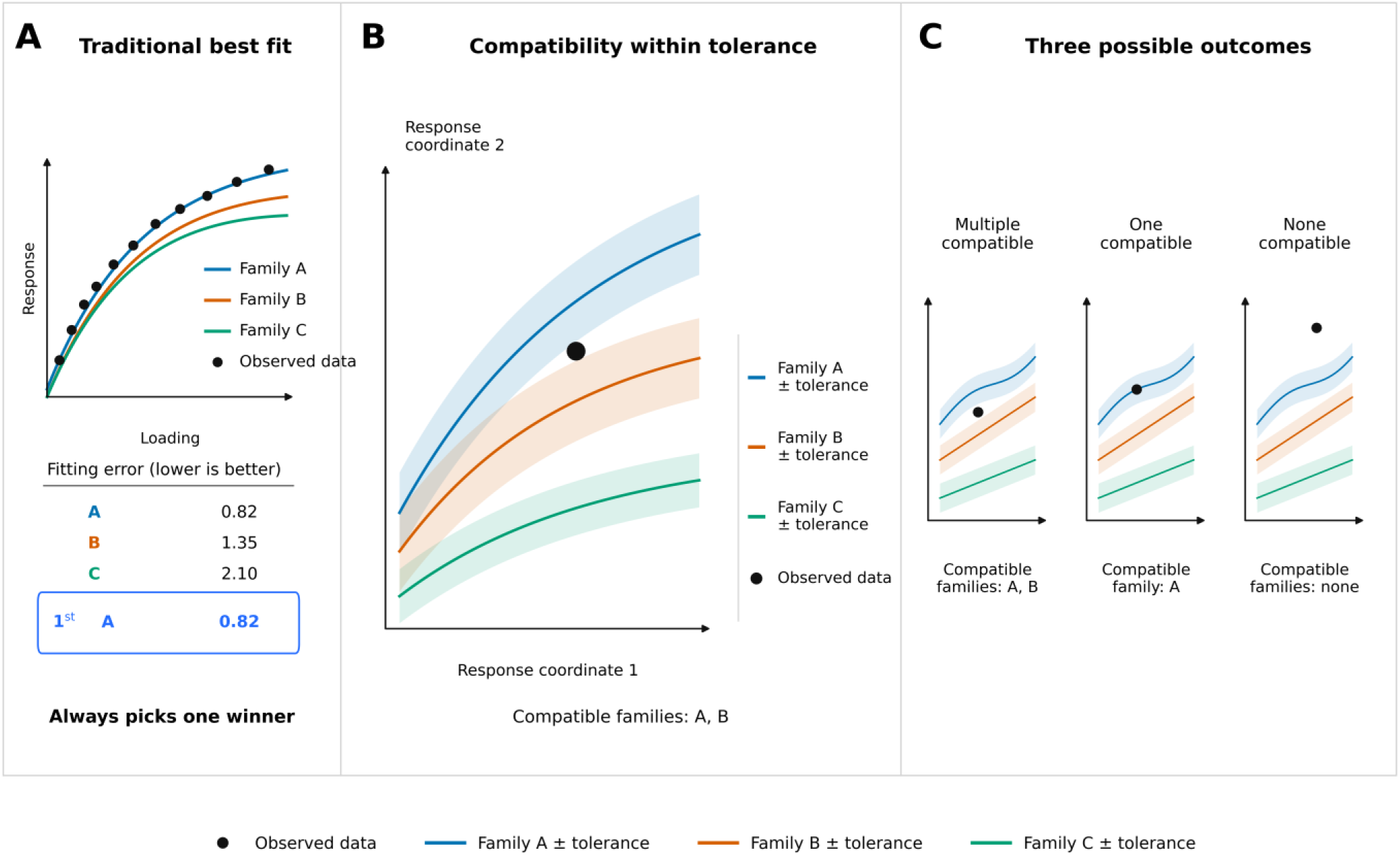
Constitutive ranking versus compatibility. (A) Relative selection returns the best candidate family even when several families are indistinguishable or all are inadequate. (B) The atlas represents each family as a response manifold surrounded by a tolerance neighborhood in a common deformation space. (C) Inference retains multiple compatible families, identifies one compatible family, or rejects the configured library. This explanatory schematic was prepared with assistance from ChatGPT (OpenAI) and reviewed by the authors; it does not depict generated research data.

## 2. Theory and computational methodology

The complete workflow is summarized in Figure 2. We define a bounded parent deformation space and a common response representation, then construct and certify the seven family manifolds. The atlas is frozen before it is queried with controlled or experimental responses. When more than one family remains, a stored witness identifies a deformation that separates their predictions.

**Figure 2.**
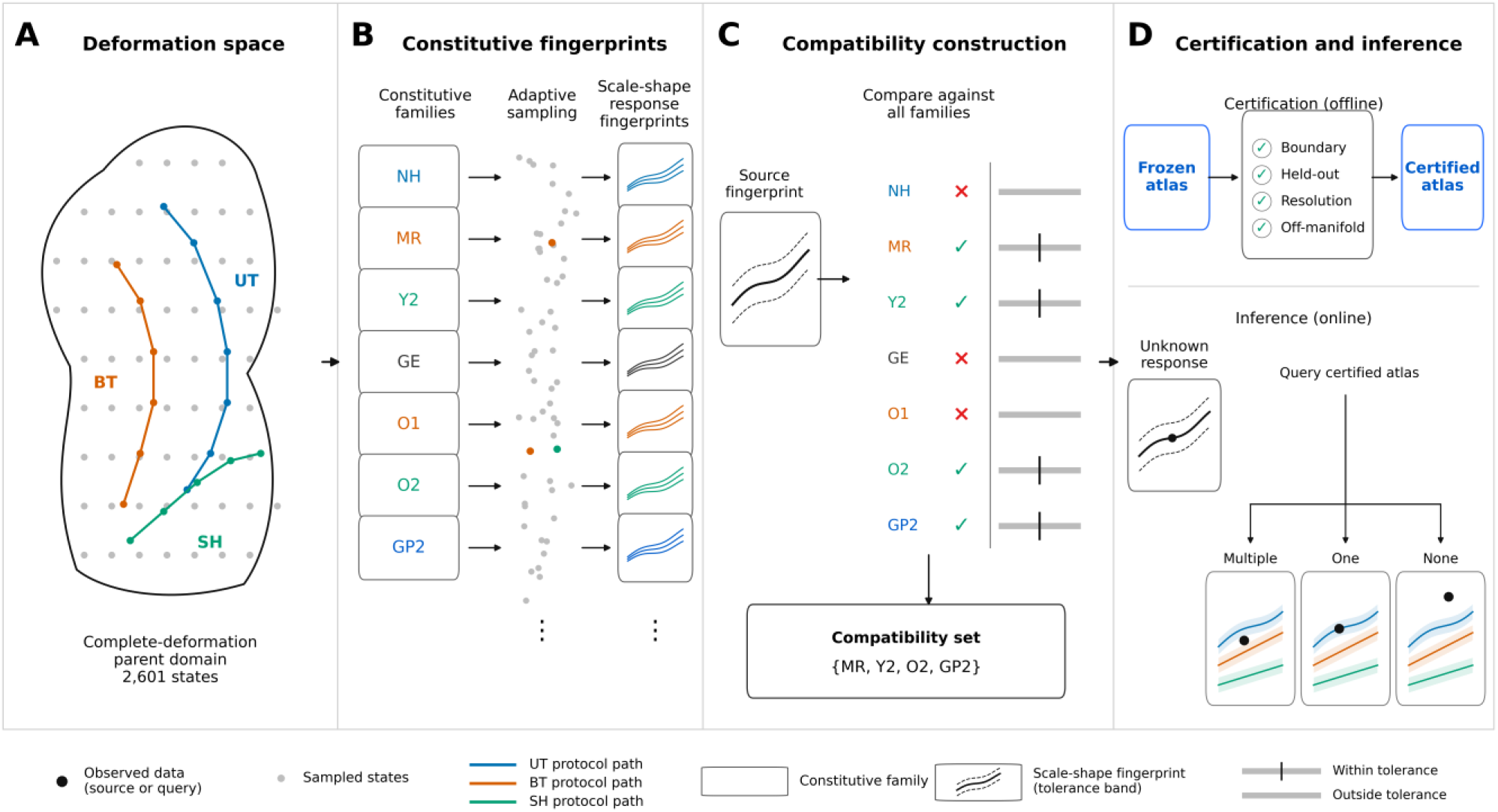
Atlas construction, certification, and inference. (A) A bounded incompressible parent domain contains the UT, BT, and SH paths. (B) Seven constitutive families are sampled in scale-shape coordinates to form response manifolds, with exact nested strata retained. (C) Every source response is fitted by every candidate family to assign a hierarchy-closed compatibility set. (D) After numerical certification, the frozen atlas is queried with an unknown response. UT, uniaxial loading; BT, equibiaxial loading; SH, simple shear. This explanatory schematic was prepared with assistance from ChatGPT (OpenAI) and reviewed by the authors; it does not depict generated research data. NH, Neo-Hookean; MR, Mooney-Rivlin; Y2, second-order Yeoh; GE, Gent; O1, one-term Ogden; O2, two-term Ogden; GP2, second-order general polynomial.

### 2.1 Constitutive library and parameter support

We define a constitutive family as a strain-energy form together with its admissible parameter domain. One parameter choice defines a constitutive instance; calibration to a target response gives a fitted instance. The candidate library is the collection of families under consideration. A family is compatible if at least one admissible instance satisfies the declared tolerance.

The library comprised Neo-Hookean (NH), Mooney-Rivlin (MR), second-order Yeoh (Y2), Gent (GE), one-term Ogden (O1), two-term Ogden (O2), and second-order general polynomial (GP2) families. Classical incompressible formulations were implemented in the invariant-based framework [20–24]. For principal stretches satisfying the incompressibility assumption (λ_1_λ_2_λ_3_ = 1), the first two invariants are

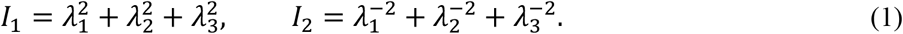

Overall stiffness and response shape were treated separately. The initial shear modulus μ_0_ sets the stress scale, whereas the remaining parameters determine the shape of the constitutive response. Table 1 gives the implemented strain-energy functions and their scale-shape mappings.

**Table 1.** Strain-energy functions and scale-shape mappings used in the constitutive library.

| Family | Strain-energy density $W$ | Initial shear modulus / scale | Shape coordinates |
| --- | --- | --- | --- |
| NH | $\frac{\mu_0}{2}(I_1 - 3)$ | $s = \mu_0/2$ | none |
| MR | $C_{10}(I_1 - 3) + C_{01}(I_2 - 3)$ | $\mu_0 = 2(C_{10} + C_{01}), s = C_{10} + C_{01}$ | $\rho = C_{01}/s; C_{10} = s(1 - \rho), C_{01} = s\rho$ |
| Y2 | $C_{10}(I_1 - 3) + C_{20}(I_1 - 3)^2$ | $\mu_0 = 2C_{10}, s = C_{10}$ | $\beta = C_{20}/C_{10}$ |
| GE | $-\frac{\mu_0 J_m}{2} \ln \left[ 1 - \frac{I_1 - 3}{J_m} \right]$ | $s = \mu_0/2$ | $\gamma = 1/J_m$ |
| O1 / O2 | $\sum_p \frac{2\mu_p}{\alpha_p^2} (\lambda_1^{\alpha_p} + \lambda_2^{\alpha_p} + \lambda_3^{\alpha_p} - 3)$ | $\mu_0 = \sum_p \mu_p, s = \mu_0/2$ | O1: $\alpha$. O2: $(w, \alpha_1, \alpha_2)$ with $\mu_1 = w\mu_0, \mu_2 = (1 - w)\mu_0$ |
| GP2 | $C_{10}(I_1 - 3) + C_{01}(I_2 - 3) + C_{20}(I_1 - 3)^2 + C_{11}(I_1 - 3)(I_2 - 3) + C_{02}(I_2 - 3)^2$ | $\mu_0 = 2(C_{10} + C_{01}), s = C_{10} + C_{01}$ | $\rho = C_{01}/s, \beta_{20} = C_{20}/s, \beta_{11} = C_{11}/s, \beta_{02} = C_{02}/s$ |

For the signed Ogden parameterization, the removable limit as an exponent approached zero was evaluated analytically. Gent parameter states that reached the locking singularity anywhere in the parent deformation domain were excluded. We assembled the construction envelopes from reported brain, adipose, breast, and meningeal properties [6,8,9,25–28] and widened them with conservative guard margins. These cross-study bounds define the computational support of the atlas. They are not probability distributions, confidence intervals, or tissue-specific priors, and every adequacy result is conditional on them. The resulting literature-core and full-support ranges are summarized in Table 2.

**Table 2.** Cross-study construction envelopes for the constitutive atlas. The literature core summarizes reported biological regimes, and the full support adds conservative guard margins.

| Family | Parameter | Literature core | Atlas support |
| --- | --- | --- | --- |
| All families | $\mu_0$ (kPa) | 0.1-12,000 | 0.05-20,000 |
| MR | $\rho$ | 0-1 | 0-1 |
| Y2 | $\beta = C_{20}/C_{10}$ | 0-20 | 0-25 |
| GE | $\gamma = 1/J_m$ | 0-0.35 | 0-0.42 |
| O1 | $\alpha$ | -25 to 25 | -30 to 30 |
| O2 | $\alpha_1, \alpha_2$ | -25 to 25 | -30 to 30 |
| O2 | $w$ | 0-1 | 0-1 |
| GP2 | $\rho$ | 0-1 | 0-1 |
| GP2 | $\beta_{20}$ | 0-20 | 0-25 |
| GP2 | $\beta_{11}$ | 0-12 | 0-15 |
| GP2 | $\beta_{02}$ | 0-20 | 0-25 |

All families used the same initial shear-modulus ranges: 0.1 ≤ μ_0_ ≤ 12,000 kPa in the literature core and 0.05 ≤ μ_0_ ≤ 20,000 kPa in the full atlas support. Because the interval spans several orders of magnitude, μ_0_ was sampled logarithmically.

### 2.2 Deformation domain and response fingerprints

Rigid rotation and permutation of the principal axes do not change the response of an isotropic material, and incompressibility requires *λ*_1_*λ*_2_*λ*_3_ = 1. An admissible state is therefore described by two independent principal stretches, or equivalently by ordered trace-free principal logarithmic strains. The parent domain was bounded by

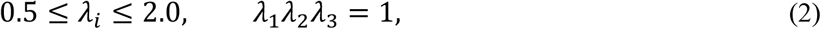

After permutation-equivalent states were removed, the intrinsic two-dimensional domain contained 2,601 states. At each state, we retained two independent pressure-free differences of principal Kirchhoff stress,

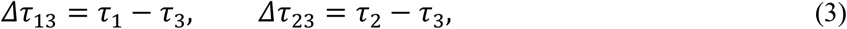

The stress differences eliminate the indeterminate hydrostatic pressure. Concatenating both components over the 2,601 frozen parent-domain deformation states produces a 5,202-component response fingerprint.

Each embedded standard protocol contained 81 states: UT over 0.70 ≤ λ ≤ 1.50, BT over 0.80 ≤ λ ≤ 1.25, and SH over -0.50 ≤ γ ≤ 0.50. Every protocol state was represented exactly in the parent set.

For material state θ, the response over the complete parent domain is

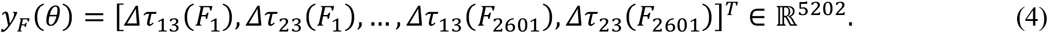

A standard protocol q *∈* {UT, BT, SH} is a deterministic projection of the same parent response,

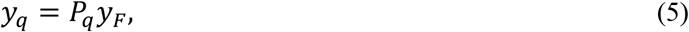

Here, *P*_*q*_ maps the ordered principal-response representation to the scalar stress observed in protocol q. UT uses (*λ*_1_, *λ*_2_, *λ*_3_) = (*λ, λ*^−1/2^, *λ*^−1/2^) and retains axial Kirchhoff stress under zero transverse traction. BT uses (*λ*_1_, *λ*_2_, *λ*_3_) = (*λ, λ, λ*^−2^) and retains in-plane Kirchhoff stress under zero thickness traction. SH uses *F* = *I* + *γ e*_1_ *⊗ e*_2_ and retains the pressure-independent shear stress *τ*_12_. In the parent domain, SH is represented by principal stretches

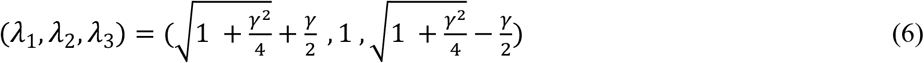

Each standard loading protocol contributes 81 scalar components, and combined views concatenate the corresponding projections from the parent general input deformation space (UT+BT, UT+SH, or BT+SH). Compatibility labels are always determined in the full 5,202-dimensional parent response space, while PCA and protocol projections are used only to visualize how much of that complete response is captured by conventional tests. Compatibility itself is always decided in the common parent space.

### 2.3 Compatibility metrics and family hierarchy

Near-zero stress components can dominate a purely relative-error measure. To avoid this behavior, each component is normalized by a local stress scale. For deformation state q and pressure-free component k, the per-unit tolerance scale for response r is

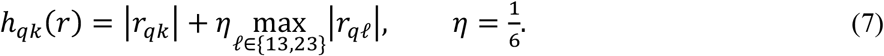

We use two metrics because atlas construction and experimental inference ask different questions. During construction, a symmetric distance compares two constitutive responses without treating either one as the observation:

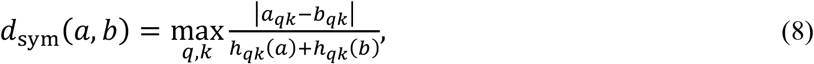

During atlas construction, the symmetric distance from a source response to a candidate family is minimized over all admissible instances of that family. This distance defines the source-atlas compatibility regions and, in the controlled examples, the minimum family-to-family noise required for overlap. Experimental inference is handled differently because a measured response does not provide its own constitutive noise scale. The observation score is therefore centered on the candidate response:

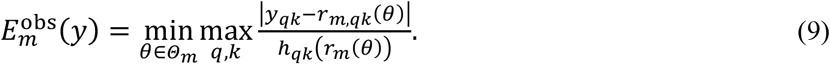

Family m is compatible when 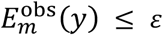. We use *ε* = 0.03 and report the normalized score 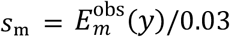 denotes compatibility. The 3% value is a declared analysis tolerance, not a universal constitutive constant.

Several families contain lower-order families exactly. Direct containments include NH within MR, Y2, GE, and O1; O1 within O2; MR within O2 and GP2; and Y2 within GP2. Transitive relations follow from these embeddings, which are listed in Supplementary Table S2. Compatibility sets are closed under this hierarchy.

A multi-family label may reflect exact mathematical non-uniqueness, not failed optimization. We therefore report the full hierarchy-closed set as the primary result. The least-complex adequate family is a secondary diagnostic. Table 3 summarizes how the comparison, compatibility, calibration, and diagnostic quantities are used.

**Table 3.** Reader’s guide to the comparison, compatibility, calibration, and diagnostic quantities used in the study.

| Quantity | Question answered | Where used | Decision meaning |
| --- | --- | --- | --- |
| Symmetric distance | How close are two constitutive responses? | Atlas construction; controlled examples | Percent noise needed for overlap |
| Observation score | Can a family explain a measured response? | Compatibility inference | Accepted when $E_m^{\text{obs}}(y) \leq \varepsilon$ |
| Normalized score, $s_m$ | How close is a fit to the tolerance boundary? | Operational reporting | $s_m \leq 1$ is compatible |
| NRMSE | How well does a fitted curve predict calibration or holdout data? | External examples | Fit error only; not an adequacy test |
| $\Delta\text{BIC}$ and $\Delta\text{AICc}$ | Which candidates survive the training-only screen? | Treloar and brain calibration | Both $\leq 2$ for retention |
| Projection distance | Does an experimental fit map near a frozen same-family state? | Frozen-atlas lookup | Must be within 3% |
| Critical-noise distance | Can a measured deformation separate two candidates? | Witness selection | >3% indicates separability |
| PCA / balanced accuracy / macro-F1 | How visible are frozen labels after protocol projection? | Diagnostic visualization | Never assigns compatibility |

### 2.4 Atlas construction and constitutive-family inference

Sampling density was adapted to constitutive sensitivity measured by the profiled Fisher metric. For MR, Y2, GE, and O1, the one-dimensional shape coordinates were distributed approximately uniformly in Fisher arc length. O2 and GP2 used scrambled Sobol pilot sets [29], followed by Fisher-volume-weighted maximin selection. Exact nested strata, parameter boundaries, and literature anchors were retained explicitly [15]. The parameter support, deformation domain, source states, and pairwise witnesses were then frozen before certification and experimental analysis.

Stiffness and shape were sampled independently. NH used a logarithmic stiffness grid, while non-NH source nodes received deterministic stratified logarithmic stiffness values. A full Cartesian product was unnecessary because the responses are homogeneous in stiffness scale and the compatibility metric is scale invariant. Each source state was then fitted to every alternative family in the 5,202-dimensional parent space, with exact nesting imposed analytically. Supplementary Table S3 gives the optimization settings.

For an unknown parent-domain response y, each candidate family is fitted independently over its admissible scale-shape domain using the candidate-centered score in Eq. (9). The preliminary compatible set is

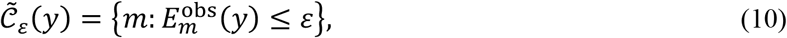

The reported set is obtained by closing this preliminary set under the known constitutive-family hierarchy:

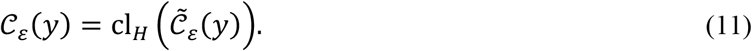

If |*C*_*ε*_| > 1, all adequate families are retained. If |*C*_*ε*_| = 1, that family is uniquely adequate only within the declared library, parameter domain, deformation domain, and tolerance. If |*C*_*ε*_| = 0, no configured family is adequate; the numerically closest family may be reported diagnostically but is not accepted as adequate. Fits close to the threshold are re-evaluated with denser target banks and multiple starting points before a label is finalized.

### 2.5 Validation and external experiments

Before it was used for inference, the frozen atlas had to pass tests of exact nesting and protocol embedding, held-out source-family containment, deformation-resolution stability, off-manifold rejection, and tolerance-boundary behavior. Supplementary Note 1 and Tables S4 and S8 report the test definitions, acceptance criteria, convergence settings, perturbation cases, and numerical outcomes.

We also asked how much of the complete-space label structure survived projection onto ALL, UT, BT, SH, UT+BT, UT+SH, and BT+SH views. After stiffness normalization, each view was represented by its first three principal components. Exact duplicate scale-free states were removed. Compatibility sets containing fewer than five remaining states were retained in the PCA plots but excluded from classifier evaluation because they could not support stratified fivefold splitting; these rare sets contained five source states in total. For each view, the PCA coordinates and their standardization were fixed before five repeats of stratified fivefold cross-validation were applied to a distance-weighted seven-nearest-neighbor classifier. Cross-validation therefore assessed label visibility within a fixed embedding rather than out-of-sample construction of the embedding. The complete-space labels remained fixed; Supplementary Note 1 gives the evaluation coverage and remaining classifier settings.

External validation used Treloar rubber data [30,31] and the human-brain data of Budday et al. [18]. For Treloar, all seven families were calibrated to uniaxial tension; planar and equibiaxial tension were withheld. A training-only information-criterion rule retained two families. We projected this pair into the frozen atlas and restricted its stored witness to deformation states present in the holdouts before examining the corresponding stresses.

Cortex, basal ganglia, corona radiata, and corpus callosum were analyzed separately. Compression was used for calibration, with tension and simple shear withheld. Discrimination required two surviving candidates, fitted states within 3% of same-family atlas nodes, and a certified source-to-target distance above 3%. For an eligible pair, the frozen witness selected the measured state before the withheld response was revealed; the other regions were not tested.

## 3. Results

### 3.1 Geometry of the constitutive-response atlas

At 3% tolerance, the seven constitutive families formed 14 hierarchy-closed compatibility sets. The frozen atlas contained 2,631 source states, each represented by a 5,202-component response over the parent domain. The most common sets were GP2 alone (29.7%) and O2 alone (21.9%); the full distribution appears in Supplementary Table S1. Because sampling was adaptive and family counts differed, these frequencies describe the frozen sample rather than unbiased manifold volumes or biological probabilities. For visualization, scale-free PCA captured 90.3% of variance in PC1-PC2 and 95.1% in PC1-PC3; all compatibility calculations remained in the original response space (Figure 3).

**Figure 3.**
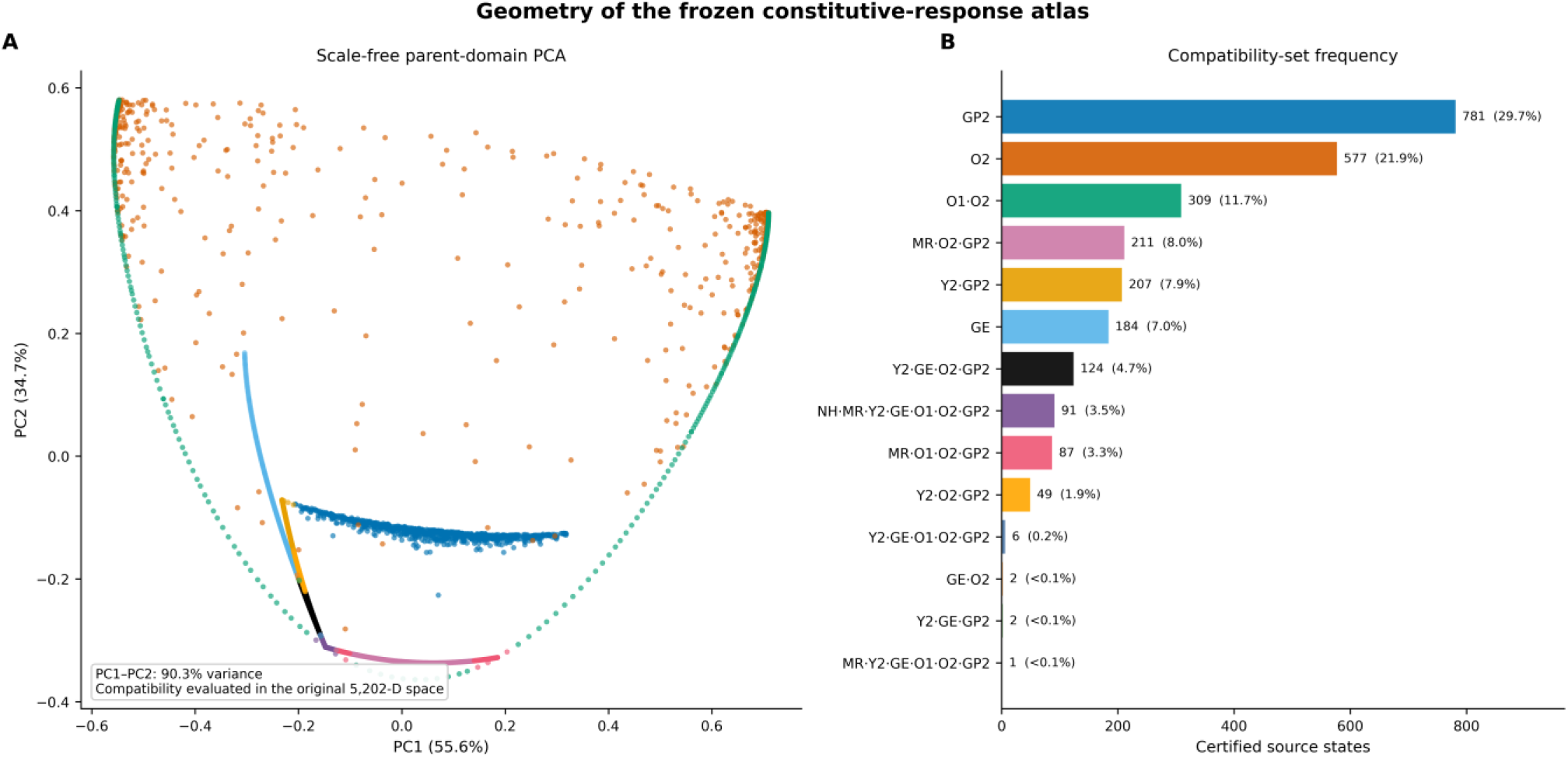
Geometry of the frozen constitutive-response atlas. (A) Scale-free PCA view of the 5,202-dimensional fingerprints, colored by compatibility set; PCA is used only for visualization. (B) Frequencies of the 14 hierarchy-closed sets across 2,631 adaptively sampled source states. Frequencies describe the frozen sample, not physical probabilities or unbiased manifold volumes. Colors assigned to compatibility sets are consistent across panels A and B. PCA, principal component analysis. NH, Neo-Hookean; MR, Mooney-Rivlin; Y2, second-order Yeoh; GE, Gent; O1, one-term Ogden; O2, two-term Ogden; GP2, second-order general polynomial.

The frozen atlas passed all seven prespecified certification criteria (Supplementary Tables S4 and S8). It recovered exact nesting and the embedded protocols, retained the generating families in clean and margin-separated held-out cases, remained stable to deformation resolution, and rejected margin-separated off-manifold responses. Recovery decreased for tests placed exactly on the high-dimensional tolerance boundary, even though the exact generating response remained feasible. Supplementary Note 1 reports the complete audit and the solver-boundary results.

### 3.2 Standard protocols as incomplete projections

A single standard protocol retains only 81 scalar stress components from the 5,202-component parent fingerprint. This projection reduced separation among the fixed complete-space labels. UT and BT preserved much of the global structure, whereas SH produced the strongest collapse: PC1 explained 95.0% of SH variance, compared with 55.6% in the parent view. Mean balanced accuracy for the evaluated fixed labels was 0.921 in the parent view, 0.879 for UT, 0.879 for BT, and 0.651 for SH. These values measure label visibility within the fixed PCA embeddings; no family was refitted within an individual protocol (Fig. 4).

**Figure 4.**
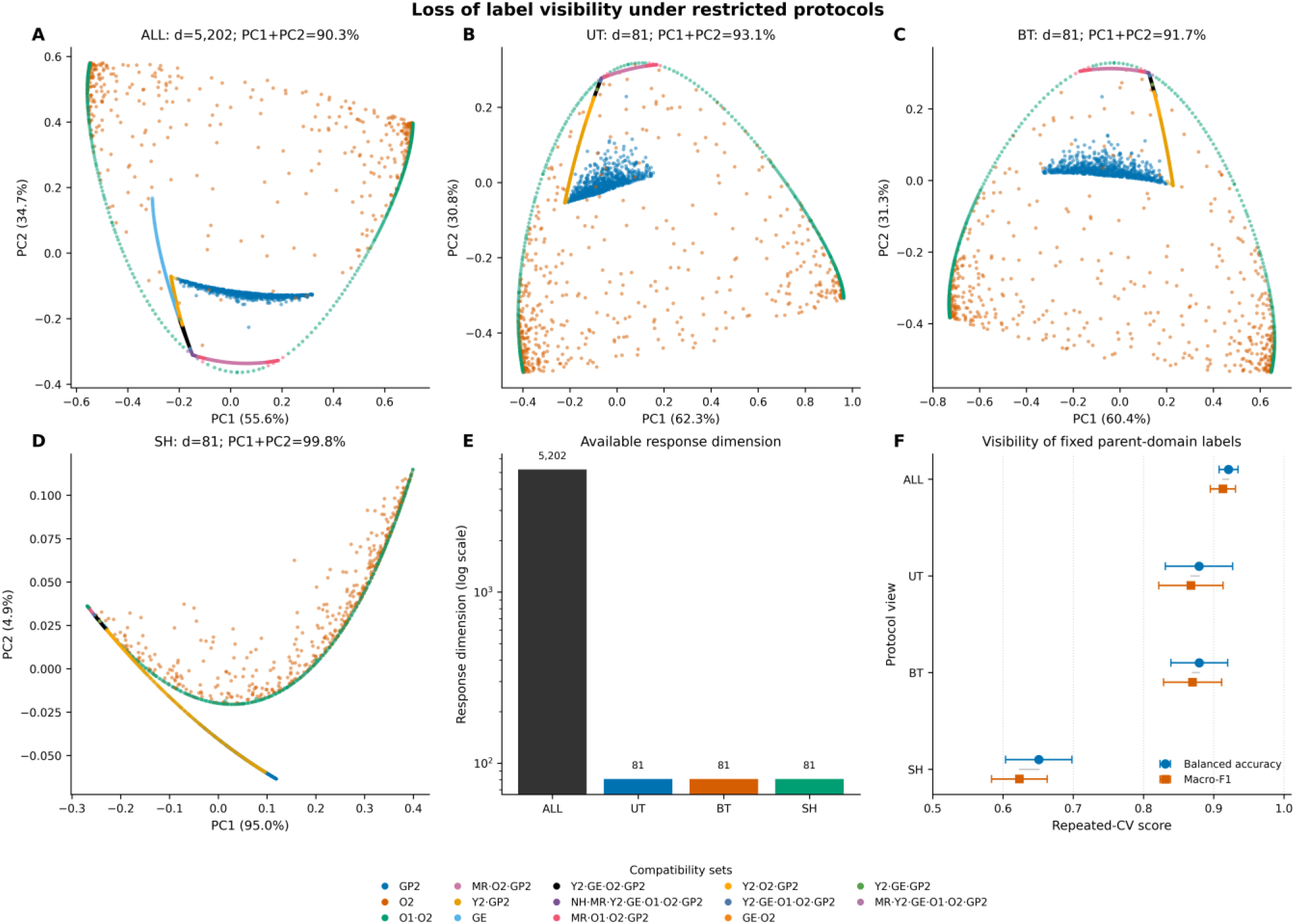
Loss of label visibility under restricted protocols. (A-D) Scale-free PCA views of the fixed complete-space labels for ALL, UT, BT, and SH. (E) Response dimensions for the parent and single-protocol views. (F) Mean balanced accuracy and macro-F1 from five repeats of stratified fivefold cross-validation; error bars indicate ±1 sample standard deviation across the 25 test-fold scores. No protocol-specific constitutive refitting was performed. ALL, complete parent-domain response; UT, uniaxial loading; BT, equibiaxial loading; SH, simple shear; PCA, principal component analysis. NH, Neo-Hookean; MR, Mooney-Rivlin; Y2, second-order Yeoh; GE, Gent; O1, one-term Ogden; O2, two-term Ogden; GP2, second-order general polynomial.

The loss of information under conventional characterization protocols has three different consequences. The controlled examples show how they can preserve genuine ambiguity, hide constitutive complexity, or make an inadequate candidate library appear successful (Fig. 5).

**Figure 5.**
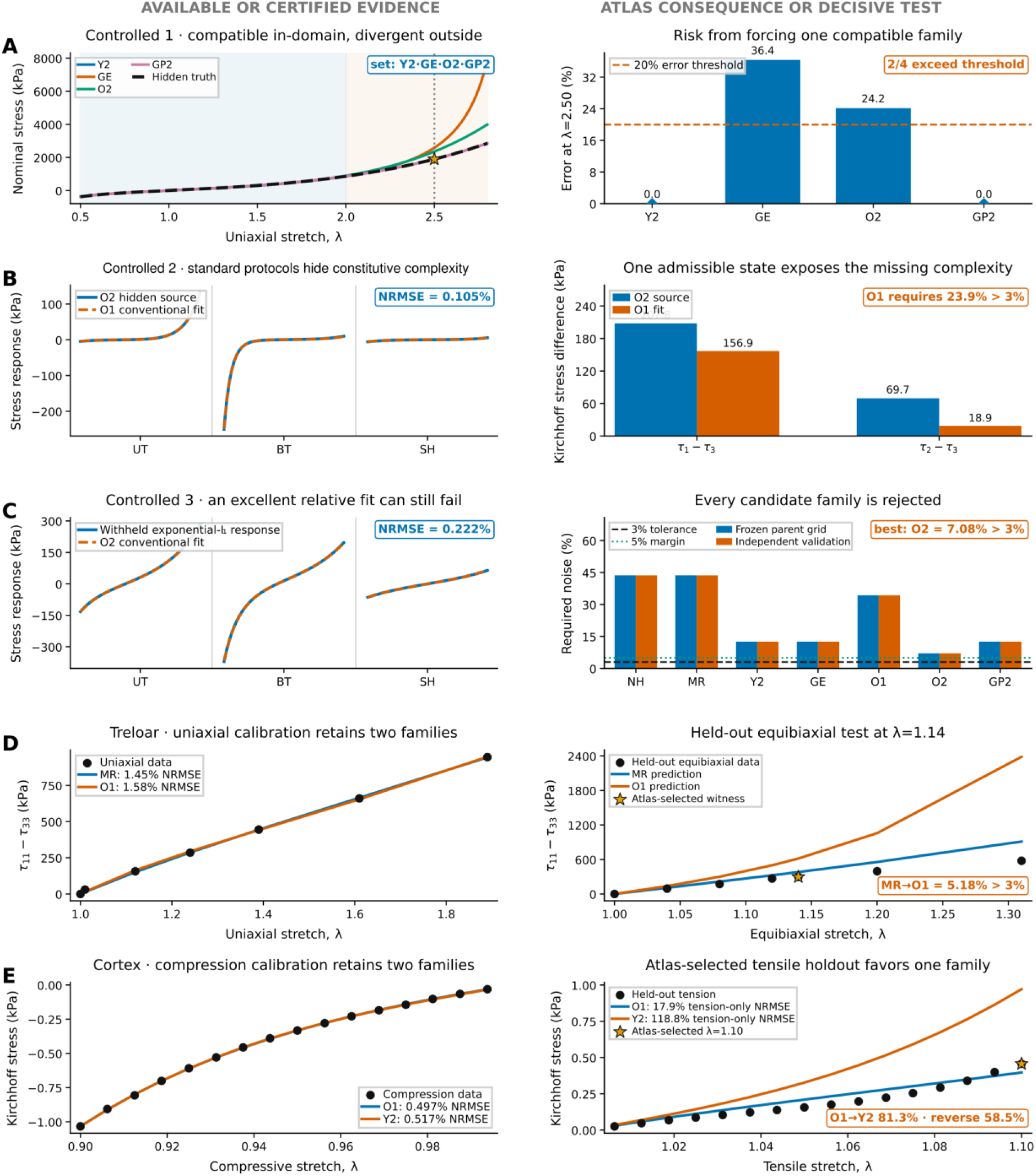
Consolidated atlas outcomes from three controlled examples and two external validations. The left column shows the response evidence available from the stated protocol or retained compatible set, and the right column shows the atlas consequence or prospective decisive test. (A) Four families are compatible with the hidden Y2 response inside the certified domain, but GE and O2 exceed the prespecified 20% extrapolation-error threshold at *λ* = 2.50. (B) O1 reproduces the standard responses of hidden O2 with 0.105% NRMSE, whereas one admissible parent-domain state exposes the missing complexity. (C) O2 fits the withheld exponential-I_1_ response on the standard protocols with 0.222% NRMSE, yet all seven families are rejected because the best full-domain score is 7.08%, above the 3% tolerance. (D) Treloar uniaxial calibration retains MR and O1; the atlas-selected equibiaxial holdout at *λ* = 1.14 favors MR within that pair. (E) Cortex compression retains O1 and Y2; the atlas-selected tensile holdout at *λ* = 1.10 favors O1 within that pair. Expanded diagnostics are provided in Supplementary Figs. S1–S5. NRMSE, normalized root-mean-square error; MR, Mooney-Rivlin; O1, one-term Ogden; O2, two-term Ogden; Y2, second-order Yeoh; GE, Gent. NH, Neo-Hookean; GP2, second-order general polynomial; UT, uniaxial loading; BT, equibiaxial loading; SH, simple shear.

### 3.3 Example 1. Preserve legitimate constitutive uncertainty

The hidden source was a second-order Yeoh instance (Y2). It was compatible with Y2, GE, O2, and GP2 throughout the certified domain 0.5 ≤ *λ*_*i*_ ≤ 2.0. We then propagated these four fitted responses to the uniaxial stretch *λ* = 2.50, selected before examining their separation. Their predictions spanned 36.4%; GE differed from the hidden Y2 response by 36.4%, and O2 differed by 24.2%. Two of the four compatible families therefore exceeded the prespecified 20% error threshold. This extrapolation is not atlas-certified. Its purpose is to show that retaining the in-domain compatibility set exposes uncertainty that arbitrarily selecting one family would conceal. The retained set and its extrapolation risk are summarized in Fig. 5A, with the expanded diagnostics in Supplementary Fig. S1.

### 3.3 Example 2. Identify necessary constitutive complexity

Standard UT, BT, and SH responses from a hidden two-term Ogden instance (O2) were reproduced by a fitted instance of the simpler one-term Ogden family (O1) with only 0.105% pooled NRMSE. Across the full parent domain, however, O1 required 23.9% symmetric noise to explain the O2 response, leaving O2 as the only compatible family at 3%. The largest in-domain separation occurred at (*λ*_1_, *λ*_2_, *λ*_3_) = (1.516, 1.320, 0.500), where the predictions differed by 32.5%. Thus, standard loading paths can conceal constitutive complexity that an admissible multiaxial state reveals. The restricted-path fit and the decisive parent-domain state are summarized in Fig. 5B, with the expanded diagnostics in Supplementary Fig. S2.

### 3.4 Example 3. Reject an inadequate candidate library

A smooth incompressible exponential −*I*_1_ law was deliberately omitted from the seven-family atlas to test candidate-library misspecification. Despite this omission, a fitted O2 instance reproduced its pooled noiseless UT, BT, and SH responses with 0.222% NRMSE.

No atlas family was adequate over the full parent domain. Even the closest family, O2, required approximately 7.08% symmetric noise in both continuous optimization and independent dense-grid validation, exceeding the 3% tolerance. The global failure witness was (*λ*_1_, *λ*_2_, *λ*_3_) = (2.0, 1.0, 0.5), with an interior witness near (1.741, 0.758, 0.758). The empty compatibility set distinguishes an excellent relative fit on standard paths from absolute adequacy over the declared domain. The standard-path fit and full-library rejection are summarized in Fig. 5C, with the expanded diagnostics in Supplementary Fig. S3.

### 3.6 External validation in Treloar rubber

Uniaxial calibration retained the Mooney-Rivlin (MR) and one-term Ogden (O1) families. Their fits were similar: MR gave 1.45% NRMSE with ΔBIC = ΔAICc = 0, and O1 gave 1.58% NRMSE with ΔBIC = ΔAICc = 1.14. Higher-dimensional O2 and GP2 fits achieved slightly lower raw errors but were excluded by AICc. Both retained fits projected within 3% of frozen same-family nodes. The nearest MR node excluded O1 at a critical-noise value of 5.18%, and all 39 MR nodes within 3% of the experimental fit gave the same exclusion. The pair and direction were fixed using uniaxial data alone.

When the stored MR-to-O1 witness was restricted to measured held-out states, it selected equibiaxial stretch *λ*_1_ = *λ*_2_ = 1.14 before the corresponding stress was examined. The measured pressure-free stress was 296.4 kPa, compared with predictions of 381.1 kPa for MR and 616.1 kPa for O1. The full planar-plus-equibiaxial holdout gave 34.0% pooled NRMSE for MR and 198.6% for O1. The atlas therefore identified a deformation that discriminated between the calibration-retained candidates. The remaining MR error means that this result validates discrimination within the pair, not global adequacy of MR for Treloar rubber. The calibration and prospective held-out test are summarized in Fig. 5D, with the frozen-atlas lookup in Supplementary Fig. S4.

### 3.7 External validation in human brain tissue

The compression-only screen was applied uniformly to cortex, basal ganglia, corona radiata, and corpus callosum [18]. Only cortex retained more than one family under the joint ΔBIC ≤ 2 and ΔAICc ≤ 2 rule. For cortex, the one-term Ogden (O1) family gave 0.497% NRMSE and the second-order Yeoh (Y2) family gave 0.517%, with their compression responses separated by only 0.696% in the atlas metric. The fitted states projected within 1.76% and 0.299% of frozen O1 and Y2 nodes, respectively. Their certified directed distances were 81.3% and 58.5%, and neighboring nodes gave the same conclusion. Restricting the stored O1-to-Y2 witness to measured holdouts selected uniaxial tension at *λ* = 1.10 before the stress was examined. At *λ* = 1.10, the measured pressure-free stress was 0.457 kPa, compared with 0.397 kPa for O1 and 0.971 kPa for Y2. Across all withheld tension and shear data, pooled NRMSE was 15.7% for O1 and 89.1% for Y2. The atlas thus resolved the compression-induced ambiguity within the retained pair, without establishing unique global adequacy of O1 for human cortex. The calibration and prospective tensile test are summarized in Fig. 5E, with the frozen-atlas scores and withheld shear response in Supplementary Fig. S5.

## 4. Discussion

### 4.1 From family ranking to constitutive adequacy

Information criteria and related methods rank fitted families within a specified library [1,3,5]. That ranking does not show whether the leading family remains adequate over a broader deformation domain. Compatibility tests this second question against the stated discrepancy tolerance. In Example 3, O2 gave the best standard-path fit, but the full-domain compatibility set was empty. Multiple compatible families are not necessarily a failure. The loading paths may be insufficient, two family manifolds may nearly coincide, or one family may be embedded in another [14,15]. In Example 1, four families were compatible inside the certified domain and separated only in extrapolation. Choosing one would assign certainty that the data did not support.

### 4.2 Loading path and candidate-library design

Which deformation is measured determines which constitutive differences are visible [7,18,19]. In Example 2, O1 reproduced the standard protocols almost exactly but failed at a multiaxial state inside the atlas. The atlas identifies separating states; optimal experimental design can then select a feasible one [32]. An empty compatibility set exposes insufficiency in the candidate library. The least-bad candidate is not relabeled as adequate. Example 3 applies bounded-error invalidation [16,17] to several nonlinear hyperelastic families and rejects the configured library. The withheld exponential−*I*_1_ law is a controlled test of that decision, not a proposed biological constitutive law.

### 4.3 Choosing and interpreting the tolerance

The 3% threshold specifies the discrepancy accepted in this analysis; it is not a universal constitutive constant. Increasing the threshold expands the compatibility sets, while decreasing it contracts them. Supplementary Note 1 reports the full persistence sweep. For a fixed deformation domain, normalization, parameter support, and tolerance, the resulting set is deterministic. It should not be interpreted as a Bayesian credible set or as a posterior probability assigned to a family. Distance from the operating threshold gives the clearest measure of robustness.

The Treloar MR-to-O1 distance was 5.18%. O1 remained excluded at 2%, 3%, and 5%, although the last case lay close to the boundary. The cortex O1-to-Y2 distance was 81.3%, showing a much wider separation. These results support robustness near the declared tolerance, not for every possible threshold.

### 4.4 What the external validation establishes

The external tests were designed to evaluate the decision process, not to establish a universally correct material law. In both datasets, a restricted calibration protocol left two plausible families. The frozen atlas then selected a measured, withheld deformation before its stress was examined. The holdouts favored MR over O1 for Treloar rubber and O1 over Y2 for cortex. These results establish prospective discrimination within each retained pair and within the declared parameter support. They do not establish unique constitutive truth.

### 4.5 Limitations and future directions

The present atlas is deliberately restricted to homogeneous, isotropic, incompressible hyperelastic behavior, seven predefined families, and bounded parameter and deformation domains. Its adequacy statements remain conditional on those supports, the response normalization, and the chosen tolerance. The frequencies reported from the adaptive sample describe that frozen sample; they are not manifold volumes or biological probabilities.

The parent domain should not be read as a requirement to measure all 2,601 states. In practice, characterization can proceed from an informative subset selected adaptively. Mapping imaging or experimental observations to a complete response representation remains a separate inverse problem. The external evidence is also limited to one classical rubber dataset and one human-brain study. Only cortex met the predefined two-family eligibility rule, and the current constitutive scope excludes loading-rate effects, anisotropy, and other biological structure.

Finite numerical searches also had more difficulty with observations placed exactly on the high-dimensional max-norm boundary than with margin-separated cases. The exact generating response remained feasible, but its numerical recovery decreased. Supplementary Note 1 reports the boundary tests.

Several extensions follow directly from the present construction. The family library can include compressible, anisotropic, fiber-reinforced, viscoelastic, and other history-dependent behavior. A more substantial next step is heterogeneous inverse characterization, in which imaging-derived deformation fields map to spatially varying compatibility sets rather than one homogeneous response. Coupling the atlas with experimental design would also allow feasible loading states to be chosen from those that best separate the remaining families [32].

## 5. Conclusion

Hyperelastic characterization should not force a unique family when the available data do not support one. The constitutive-response atlas makes this principle operational over a declared deformation domain and tolerance. It retains every adequate family and returns an empty set when the configured library is insufficient. The controlled examples showed that standard loading paths may leave genuine ambiguity, hide constitutive complexity, or make an inadequate library appear successful. In the external datasets, the frozen atlas selected held-out deformations that distinguished MR from O1 in Treloar rubber and O1 from Y2 in human cortex. Family-level compatibility can therefore communicate the uncertainty that remains and guide the next measurement when a restricted protocol is not informative enough.

## Data availability

The Treloar validation data are publicly available in machine-readable form from the thermalCANN repository (https://github.com/llamm-de/thermalCANN/tree/main/data/Treloar), with the original experimental and reproduction sources given in [30,31]. The human-brain validation data are available from the HyperSmart material repository (https://github.com/dhnp90/HyperSmart/tree/main/src/material_repository), with the experimental source given in [18]. The frozen constitutive atlas, certification outputs, experimental-validation outputs, and analysis notebook are archived on Zenodo (version v1; https://doi.org/10.5281/zenodo.22818725). The code repository is available at https://github.com/aasad009/constitutive-compatibility-atlas.

## CRediT authorship contribution statement

Amirreza Asadi: Conceptualization, Methodology, Software, Validation, Formal analysis, Investigation, Data curation, Visualization, Writing – Original Draft.

Kaveh Laksari: Conceptualization, Methodology, Supervision, Validation, Formal analysis, Investigation, Resources, Data curation, Software, Writing – Review & Editing, Visualization, Project administration, Funding acquisition.

## Declaration of generative AI and AI-assisted technologies in the manuscript preparation process

During preparation of this work, the authors used ChatGPT (OpenAI; GPT-5.6 Sol) to assist with manuscript organization, language editing, formatting, and the layout and refinement of the explanatory schematics in Figures 1 and 2. The tool was not used to generate or modify research data. The authors reviewed and edited all content, verified the explanatory figures, and take full responsibility for the publication.

