## Supplementary Data for "Noise-Aware Constitutive Compatibility Atlas for Hyperelastic Material Characterization"

Supplementary Figures S1–S5, Note 1, and Tables S1–S8 provide the numerical certification record, the complete compatibility-set distribution, exact hierarchy embeddings, frozen construction settings, prespecified acceptance criteria, external-validation robustness audits, controlled-example source identifiers, and local atlas-transfer checks referenced in the main manuscript. A constitutive family combines a strain-energy form with its admissible parameter domain, whereas a state or instance is one parameter choice within that family. Family abbreviations used throughout are Neo-Hookean (NH), Mooney-Rivlin (MR), second-order Yeoh (Y2), Gent (GE), one-term Ogden (O1), two-term Ogden (O2), and second-order general polynomial (GP2). Reference numbers follow the main manuscript.

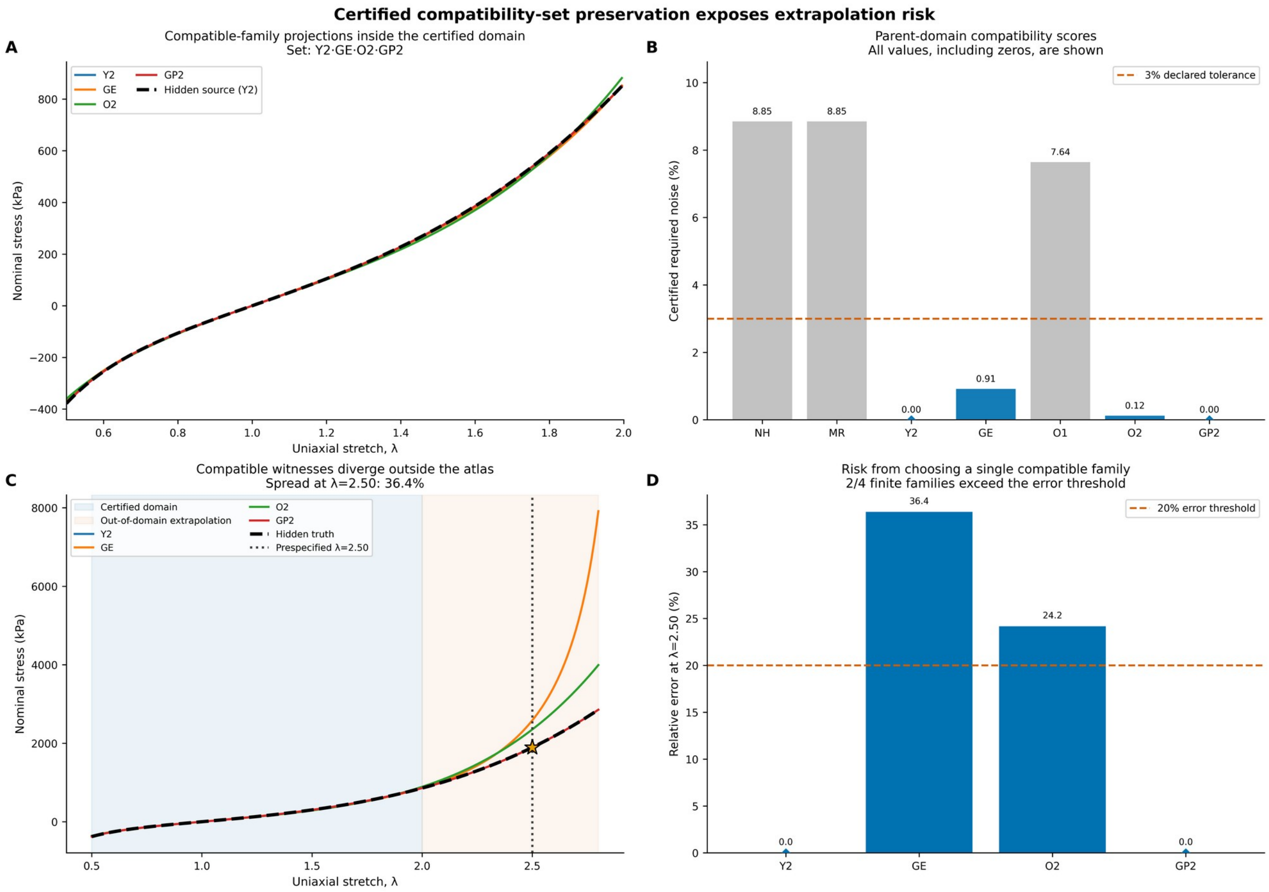


**Supplementary Figure S1.** Expanded diagnostics for controlled Example 1. (A) The retained Y2, GE, O2, and GP2 responses are visually coincident on the certified uniaxial range. (B) Parent-domain compatibility scores retain those four families at the declared 3% tolerance. (C, D) Their predictions diverge outside the certified range; GE and O2 exceed the prespecified 20% error threshold at λ = 2.50. Panels C and D are condensed in main Figure 5A.

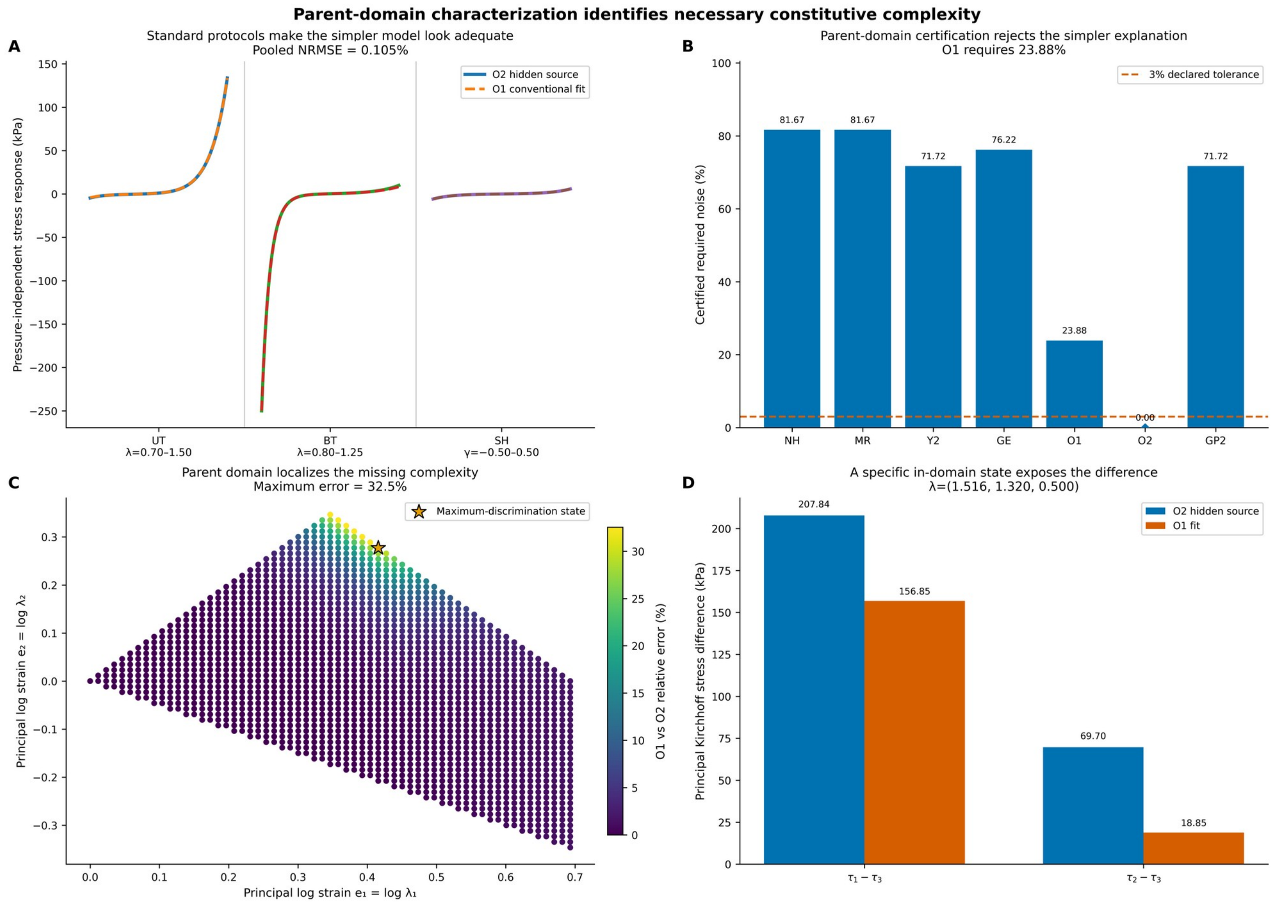


**Supplementary Figure S2.** Expanded diagnostics for controlled Example 2. (A) O1 reproduces the hidden O2 response on UT, BT, and SH with 0.105% pooled NRMSE. (B) Parent-domain certification rejects O1, which requires 23.88% noise. (C) The pointwise error map localizes the maximum-discrimination state. (D) The two independent stress differences at that state expose the missing complexity. Panels A and D are condensed in main Figure 5B.

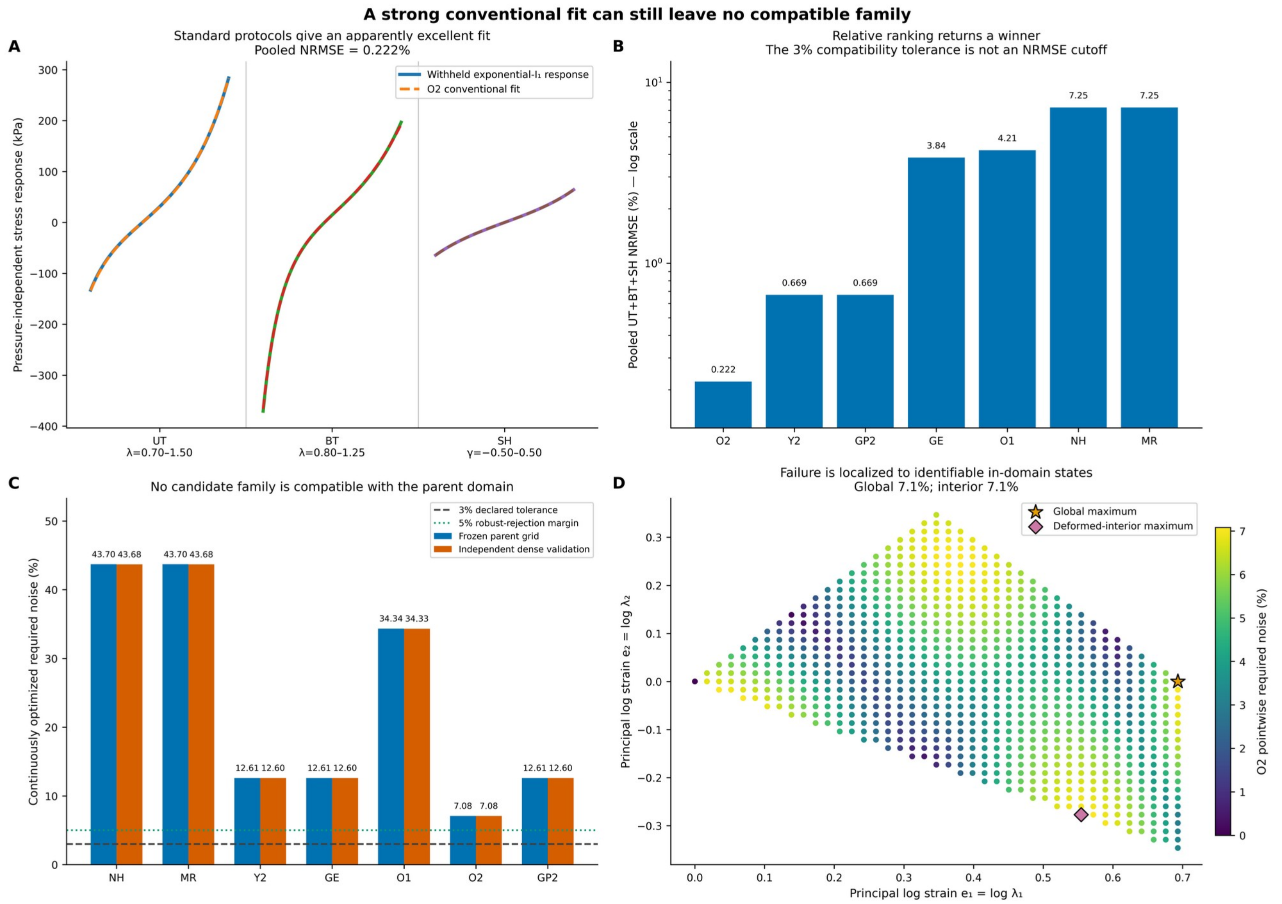


**Supplementary Figure S3.** Expanded diagnostics for controlled Example 3. (A) O2 fits the withheld exponential-$I_1$ response on UT, BT, and SH with 0.222% pooled NRMSE. (B) Relative standard-protocol ranking returns O2 as the best candidate. (C) Every atlas family remains above the 3% compatibility tolerance in the parent domain. (D) The pointwise map localizes the global and deformed-interior failure maxima. Panels A and C are condensed in main Figure 5C.

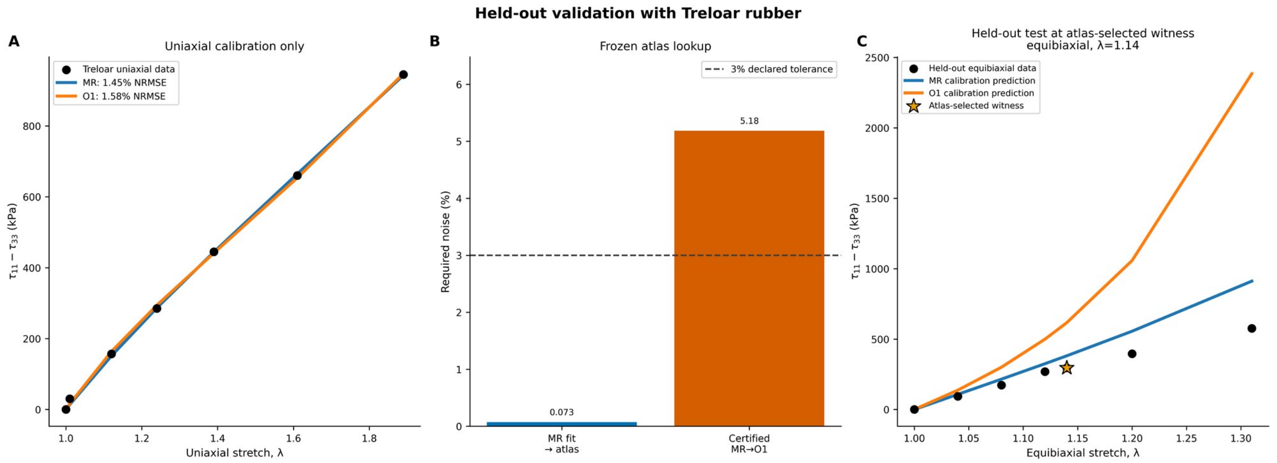


**Supplementary Figure S4.** Expanded diagnostics for the Treloar validation. (A) Uniaxial calibration retains MR and O1. (B) The fitted MR response projects within 0.073% of the frozen atlas, while the certified MR-to-O1 separation is 5.18%, above the 3% tolerance. (C) The prospectively selected equibiaxial holdout at λ = 1.14 favors MR within the retained pair. Panels A and C are condensed in main Figure 5D.

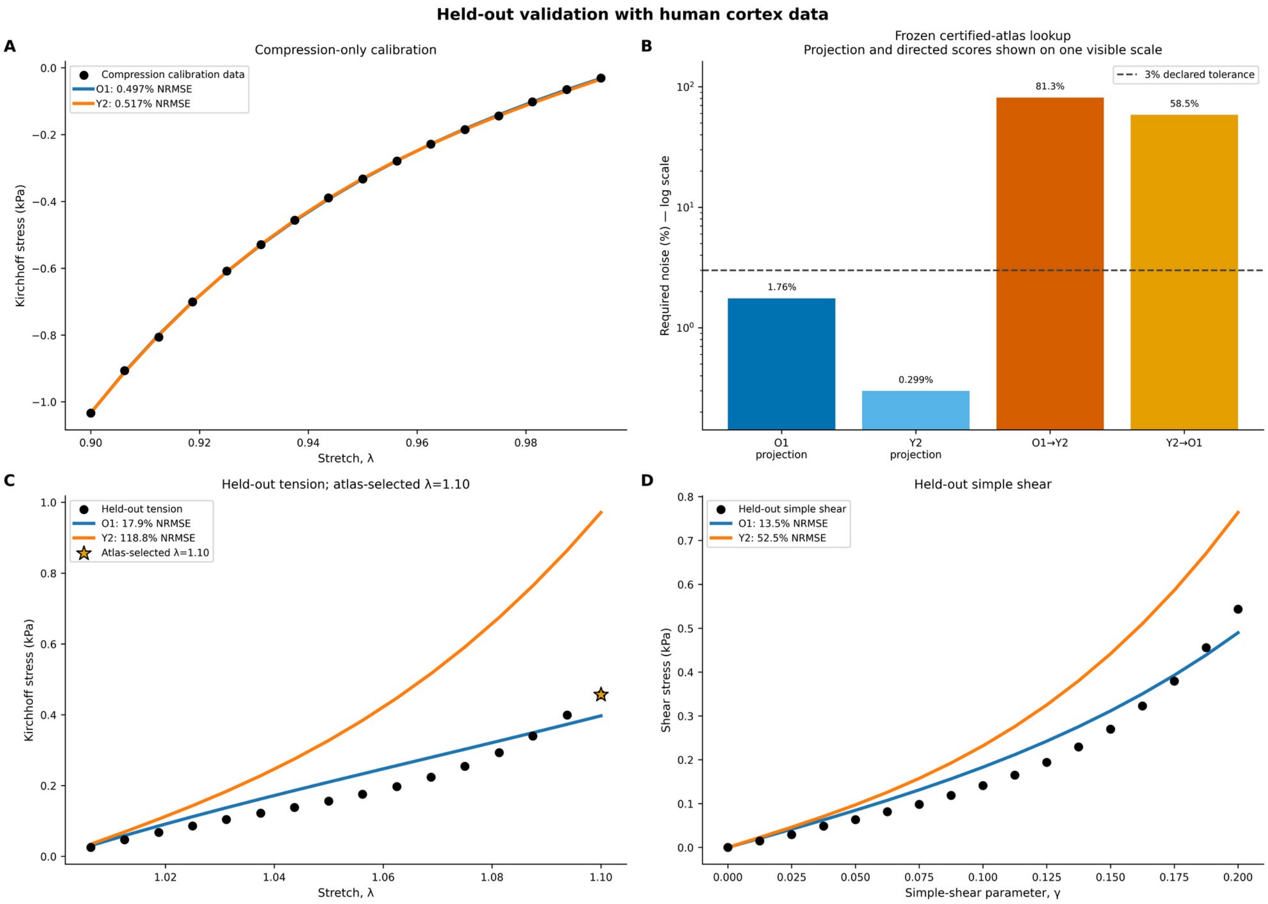


**Supplementary Figure S5.** Expanded diagnostics for the human-cortex validation. (A) Compression-only calibration retains O1 and Y2. (B) The fitted states project within 1.76% and 0.299% of frozen same-family nodes, and the certified directed distances are 81.3% from O1 to Y2 and 58.5% in the reverse direction. (C) The prospectively selected tensile holdout at λ = 1.10 favors O1. (D) The completely withheld simple-shear response gives the same within-pair preference. Panels A and C are condensed in main Figure 5E.

**Supplementary Table S1.** Complete distribution of the 14 certified hierarchy-closed compatibility sets at the 3% tolerance.

| Compatibility set | n states | % of atlas |
|---|---|---|
| GP2 | 781 | 29.68 |
| O2 | 577 | 21.93 |
| O1\|O2 | 309 | 11.74 |
| MR\|O2\|GP2 | 211 | 8.02 |
| Y2\|GP2 | 207 | 7.87 |
| GE | 184 | 6.99 |
| Y2\|GE\|O2\|GP2 | 124 | 4.71 |
| NH\|MR\|Y2\|GE\|O1\|O2\|GP2 | 91 | 3.46 |
| MR\|O1\|O2\|GP2 | 87 | 3.31 |
| Y2\|O2\|GP2 | 49 | 1.86 |
| Y2\|GE\|O1\|O2\|GP2 | 6 | 0.23 |
| GE\|O2 | 2 | 0.08 |

| Compatibility set | n states | % of atlas |
|---|---|---|
| Y2\|GE\|GP2 | 2 | 0.08 |
| MR\|Y2\|GE\|O1\|O2\|GP2 | 1 | 0.04 |

Note: Percentages summarize the composition of the adaptively sampled frozen source atlas and should not be interpreted as biological probabilities or unbiased geometric volumes of the constitutive manifolds.

**Supplementary Table S2.** Exact lower-dimensional embeddings used for hierarchy closure.

| Containment | Exact embedding in atlas coordinates |
|---|---|
| NH ⊂ MR | $\rho = 0$ |
| NH ⊂ Y2 | $\beta = 0$ |
| NH ⊂ GE | $\gamma = 1 / J_m = 0$ |
| NH ⊂ O1 | $\alpha = 2$ |
| O1 ⊂ O2 | $w = 1$ or $0$, or $\alpha_1 = \alpha_2 = \alpha$ |
| MR ⊂ O2 | $\alpha_1 = -2$, $\alpha_2 = 2$, $w = \rho$ |
| MR ⊂ GP2 | $\beta_{20} = \beta_{11} = \beta_{02} = 0$, same $\rho$ |
| Y2 ⊂ GP2 | $\rho = \beta_{11} = \beta_{02} = 0$, $\beta_{20} = \beta$ |

**Supplementary Table S3.** Frozen atlas-construction settings.

| Setting | Frozen value |
|---|---|
| Global random seed | 20260806 |
| Parent discretization | 2,601 states; 5,202 response components |
| NH source states | 25 logarithmically spaced stiffness states |
| MR / Y2 / GE / O1 | 301 Fisher-arc states per family |
| O2 | 601 Fisher-volume-weighted maximin states |
| GP2 | 801 Fisher-volume-weighted maximin states |
| Total source states | 2,631 |
| Directed alternative-family fits | 15,786 |
| Scalar Fisher pilot / target grids | 8,001 / 401 states per scalar family |
| O2 pilot / target banks | 12,288 / 2,200 states |
| GP2 pilot / target banks | 16,384 / 3,000 states |
| Near-boundary continuous refinement | 5 starts; 150 iterations / 1,800 evaluations maximum; 16 scale and 24 shape screening candidates |

**Supplementary Table S4.** Predefined numerical certification criteria.

| Criterion | Acceptance condition |
|---|---|
| Frozen-input integrity | Required frozen outputs and baseline audits internally consistent |

| Criterion | Acceptance condition |
|---|---|
| Dense boundary certification | Multi-seed re-evaluation completed for all selected near-threshold / rare cases |
| Certified source inclusion | Generating family retained for every certified source state |
| Held-out source inclusion | Minimum family/regime inclusion $\geq 0.995$ for clean and 2.7%-bounded observations |
| Held-out oracle inclusion | Exact generating-family response within the tolerance tube for every hard-validation observation |
| Parent-resolution convergence | Non-boundary agreement $\geq 0.99$ and median $\left\|s_{2601} - s_{5201}\right\| \leq 0.0015$ |
| Stratified off-manifold rejection | All selected margin-separated observations remain outside every family tolerance tube |

**Supplementary Table S5.** Supplementary direct fitted-state robustness audit across all six pairs formed by the four ΔBIC ≤ 10 uniaxial-training-plausible families. These distances are diagnostics between the specific UT-fitted parameter states and are distinct from the manuscript-primary frozen-atlas lookup.

| Pair | max ΔBIC | UT distance (%) | Full-domain distance (%) | Preselected held-out witness | Witness / pooled holdout preference |
|---|---|---|---|---|---|
| MR–O1 | 1.14 | 1.98 | 52.92 | Equibiaxial, λ=1.31 | MR / MR |
| MR–O2 | 2.34 | 0.82 | 43.02 | Equibiaxial, λ=1.31 | MR / MR |
| O1–O2 | 2.34 | 1.16 | 14.34 | Equibiaxial, λ=1.31 | O2 / O2 |
| O2–GP2 | 4.42 | 0.15 | 16.98 | Planar, λ=1.87 | GP2 / GP2 |
| MR–GP2 | 4.42 | 0.67 | 28.86 | Equibiaxial, λ=1.31 | MR / MR |
| O1–GP2 | 4.42 | 1.31 | 30.09 | Equibiaxial, λ=1.31 | GP2 / GP2 |

Note: The witness preference and pooled holdout preference agree for all six pairs. The primary manuscript result instead uses the frozen certified MR node lookup and stored MR→O1 witness; Table S5 is included only to show that the qualitative hidden-divergence conclusion is not unique to one BIC-supported fitted-state pair.

**Supplementary Table S6.** Human-brain regional pre-holdout eligibility and cortex frozen-atlas validation summary.

| Region | Strict support | Eligible? | Frozen-atlas result | Withheld validation |
|---|---|---|---|---|
| Cortex | O1 \| Y2 | Yes | Compression pair 0.70%; projections 1.76% / 0.30%; O1→Y2 81.32% | Tension λ=1.10 selected; O1 favored at witness and pooled holdout |
| Basal ganglia | Y2 | No | Single strict family | Not tested |
| Corona radiata | Y2 | No | Single strict family | Not tested |
| Corpus callosum | O1 | No | Single strict family | Not tested |

Note: The four anatomical regions are regional cases from the single Budday et al. [18] human-brain study. Eligibility was determined before tension or shear stresses were examined. A region required at least two families satisfying both ΔBIC ≤ 2 and ΔAICc ≤ 2 on compression-only training, compression-pair distance ≤ 3%, same-family frozen-atlas projections ≤ 3%, and certified source→target incompatibility > 3%. The three ineligible regions therefore did not pose a two-family ambiguity test and were not counted as atlas failures.

**Supplementary Table S7.** Post-selection local transfer stability for the two external experimental validations. The audit evaluates neighboring same-family frozen source nodes after the experimental pair has already been selected; it is not used for candidate selection or witness selection.

| Case / direction | Nearest projection (%) | Nodes within 3% | Fraction excluding target | Certified distance among local nodes (%) |
|---|---|---|---|---|
| Treloar MR→O1 | 0.073 | 39 | 39/39 (100%) | 4.31 / 5.18 / 6.61 |
| Treloar O1→MR | 0.246 | 1 | 1/1 (100%) | 22.74 / 22.74 / 22.74 |
| Cortex O1→Y2 | 1.755 | 1 | 1/1 (100%) | 81.32 / 81.32 / 81.32 |
| Cortex Y2→O1 | 0.299 | 7 | 7/7 (100%) | 53.29 / 53.97 / 62.13 |

Note: Values in the final column are minimum / median / maximum directed certified critical-noise distances among same-family nodes lying within the 3% projection neighborhood. For cortex O1→Y2, only one source node lay within the strict 3% projection neighborhood; nevertheless, all ten nearest O1 nodes excluded Y2, with directed critical-noise distances of 80.60–81.84%. All local audits were performed after the primary experimental candidates had already been frozen and therefore served only as transfer-stability checks.

### Supplementary Note 1. Numerical certification and audit details

This note contains the numerical checks that establish whether the frozen atlas can be trusted. They are kept separate from the main article because they support implementation reliability rather than the primary scientific narrative. The primary controlled-example and external-validation panels remain in the article, while the expanded diagnostics are provided in Supplementary Figures S1–S5.

#### 1.1 Frozen certification design

Certification began only after the deformation domain, parameter support, source states, and pairwise witnesses had been frozen. All 2,631 source states and 15,786 directed alternative-family fits were checked. Near-threshold, rare-region, and discontinuity cases were re-evaluated with denser target banks and independent starts. Certification could refine an existing pairwise distance, but it could not change the atlas geometry or regenerate source states.

Exact nested identities and the embedded UT, BT, and SH responses were verified against the parent-domain representation. Independent validation then used 72 held-out observations per family (504 total) under clean, 2.7%-bounded, 3%-clipped Gaussian, and 3%-clipped spatially correlated perturbations. Clean and 2.7%-bounded cases formed the hard containment set and required at least 99.5% generating-family inclusion. Exactly clipped 3% cases were treated as boundary stress tests because a max-norm perturbation can lie effectively on the acceptance boundary.

Resolution independence was assessed with 627 fixed source-target witnesses on parent grids containing 1,301, 2,601, and 5,201 states. The prespecified criterion required non-boundary label agreement above 0.99 and a median score change below 0.0015 between the two finest grids. Off-manifold rejection was tested separately with 46 margin-separated synthetic observations. Threshold persistence was evaluated at 0%, 1%, 2%, 3%, and 5% tolerance.

#### 1.2 Protocol-visibility diagnostic

Protocol visibility was evaluated for ALL, UT, BT, SH, UT+BT, UT+SH, and BT+SH views. Responses were restricted to each view and normalized for stiffness before PCA, while the final parent-domain labels remained fixed. Visibility in the first three PCs was measured by repeated stratified five-fold cross-validation with a distance-weighted seven-nearest-neighbor classifier. These calculations ask whether complete-space labels remain separable after projection; they are not protocol-specific constitutive refits.

UT and BT retained substantial global structure, whereas SH was strongly compressed. For SH, PC1 explained 94.97% of variance and PC1-PC2 explained 99.85%, compared with 55.62% and 90.34% for the complete parent response. Repeated cross-validated balanced accuracy was 0.921 for the parent, 0.879 for UT, 0.879 for BT, and 0.651 for SH; the corresponding macro-F1 values were 0.913, 0.868, 0.870, and 0.623.

#### 1.3 Certification outcomes and boundary limitation

All seven prespecified criteria were satisfied. Generating-family inclusion was 100% for all 2,631 frozen source states and all 252 clean or 2.7%-bounded held-out observations, with no oracle failures. Dense boundary certification re-evaluated 537 pairwise fits and changed 47 source labels (1.79%) without changing the 14 observed compatibility-set types. Exact constitutive nesting and the embedded protocol responses were recovered to numerical precision.

Across the 627 resolution tests, the median score change from 2,601 to 5,201 parent states was zero and non-boundary label agreement was 100%. All 46 margin-separated off-manifold observations remained outside the configured library after dense revalidation. Compatibility expanded gradually with tolerance: the mean number of compatible families was 1.54 at 0%, 1.68 at 1%, 1.76 at 2%, 1.86 at 3%, and 1.99 at 5%.

The exactly 3%-clipped cases were retained as boundary stress tests. The exact generating-family oracle remained inside the tolerance tube in every case, but numerical source-family recovery decreased to 50.8% for clipped-Gaussian perturbations and 86.5% for spatially correlated perturbations. The result reflects finite-search sensitivity when thousands of max-norm components approach the acceptance boundary. It supports margin-separated observations for hard numerical validation and is reported as a solver-boundary limitation.

#### 1.4 Controlled examples and secondary external-validation diagnostics

The controlled demonstrations used frozen Y2 source state #30 in Example 1 and frozen O2 source state #184 in Example 2. Example 3 used the withheld incompressible exponential-$I_1$ response described in the main manuscript. These identifiers connect the narrative examples to the stored atlas outputs without interrupting the main-text discussion.

For Treloar rubber, the fitted MR state projected within 0.0725% of frozen MR node #210, and all 39 MR nodes within a 3% neighborhood excluded O1. The fitted O1 state projected within 0.246% of its nearest frozen node, which excluded MR. Supplementary Figure S4B shows the lookup. The 5.18% frozen MR-to-O1 distance is a family-level minimum over admissible O1 states, whereas the larger direct distance in Table S5 compares two specific uniaxially fitted states. The difference between experimental and stored-node stiffness is not a projection error because the atlas separates constitutive shape from overall stiffness and uses a scale-invariant response metric.

For cortex, the fitted O1 and Y2 states projected within 1.76% (Supplementary Figure S5B) of frozen O1 node #84 and within 0.299% (0.30%) of frozen Y2 node #274. Their certified directed distances were 81.3% from O1 to Y2 and 58.5% in the reverse direction. These secondary checks support transfer of the frozen atlas to the fitted states; they do not change the pairwise, conditional interpretation of the external tests.

**Supplementary Table S8.** Outcomes for the seven prespecified numerical certification criteria.

| Criterion | Prespecified test | Acceptance condition | Outcome |
|---|---|---|---|
| Frozen-input integrity | Required frozen outputs and baseline audits | Internally consistent | Passed |
| Dense boundary certification | 537 selected near-threshold and rare pairwise fits | Multi-seed re-evaluation completed | Completed; 47 labels changed; 14 set types unchanged |
| Certified source inclusion | 2,631 frozen source states | Generating family retained for every certified source state | 2,631/2,631 (100%) |
| Held-out source inclusion | 252 clean and 2.7%-bounded observations | Minimum family/regime inclusion at least 99.5% | 100% in every family/regime |
| Held-out oracle inclusion | 252 hard-validation observations | Exact generating-family response inside the tolerance tube for every observation | 252/252 (100%); no oracle failures |
| Parent-resolution convergence | 627 fixed witnesses on 1,301/2,601/5,201-state grids | Non-boundary agreement at least 0.99; median $\|S2601 - S5201\|$ at most 0.0015 | 100% agreement; median change 0 |
| Stratified off-manifold rejection | 46 margin-separated observations | Remain outside every family tolerance tube | 46/46 (100%) rejected |
